# Exploring Frequency-dependent Brain Networks from ongoing EEG using Spatial ICA during music listening

**DOI:** 10.1101/509802

**Authors:** Yongjie Zhu, Chi Zhang, Petri Toiviainen, Minna Huotilainen, Klaus Mathiak, Tapani Ristaniemi, Fengyu Cong

## Abstract

Recently, exploring brain activity based on functional networks during naturalistic stimuli especially music and video represents an attractive challenge because of the low signal-to-noise ratio in collected brain data. Although most efforts focusing on exploring the listening brain have been made through functional magnetic resonance imaging (fMRI), sensor-level electro- or magnetoencephalography (EEG/MEG) technique, little is known about how neural rhythms are involved in the brain network activity under naturalistic stimuli. This study exploited cortical oscillations through analysis of ongoing EEG and musical feature during free-listening to music. We used a data-driven method that combined music information retrieval with spatial Independent Components Analysis (ICA) to probe the interplay between the spatial profiles and the spectral patterns. We projected the sensor data into cortical space using a minimum-norm estimate and applied the Short Time Fourier Transform (STFT) to obtain frequency information. Then, spatial ICA was made to extract spatial-spectral-temporal information of brain activity in source space and five long-term musical features were computationally extracted from the naturalistic stimuli. The spatial profiles of the components whose temporal courses were significantly correlated with musical feature time series were clustered to identify reproducible brain networks across the participants. Using the proposed approach, we found brain networks of musical feature processing are frequency-dependent and three plausible frequency-dependent networks were identified; the proposed method seems valuable for characterizing the large-scale frequency-dependent brain activity engaged in musical feature processing.

## Introduction

Understanding how our brain perceives complex and continuous inputs from the real-world has been an attractive problem in cognitive neuroscience in the past few decades. Brain imaging technology provides an opportunity to address this issue. However, revealing brain states is generally more difficult during real-word experiences than those recorded brain activities during resting state or simplified static stimuli like controlled and rapidly repeated stimuli (U. Hasson, Malach, & Heeger, 2010; Malcolm, Groen, & Baker, 2016; Spiers & Maguire, 2007). Some studies have tried to probe brain processes involved in a natural and realistic task. Hasson et al. demonstrated consistent and synchronized brain activity patterns across movie viewers with functional magnetic resonance imaging (fMRI) (Uri Hasson, Nir, Levy, Fuhrmann, & Malach, 2004). After that, Lankinen et al. conducted the similar experiment to examine the intersubject correlation with magnetoencephalography (MEG) (K Lankinen, J Saari, Ritta Hari, & M Koskinen, 2014; Lankinen et al., 2018). Shou et al. probed the neural activations a real-world air traffic control task (Shou, Ding, & Dasari, 2012). The question of how to disentangle stimuli-induced brain activity from spontaneous activity still remains open for scientific research due to the complexity of natural situations.

Recently, the brain state under the naturalistic stimuli including music and movie has been investigated through fMRI (V. Alluri et al., 2012; V. Alluri et al., 2013; Burunat, Alluri, Toiviainen, Numminen, & Brattico, 2014; I. Burunat et al., 2016; Toiviainen, Alluri, Brattico, Wallentin, & Vuust, 2014), MEG (Koskinen et al., 2013; K. Lankinen, J. Saari, R. Hari, & M. Koskinen, 2014) and electroencephalography (EEG) (Cong, Alluri, Nandi, Toiviainen, Fa, Abu-Jamous, Gong, Craenen, Poikonen, Huotilainen, et al., 2013; Daly et al., 2014; Daly et al., 2015; R. S. Schaefer, Desain, & Farquhar, 2013; I. Sturm, Dahne, Blankertz, & Curio, 2015). Alluri et al. explored the neural correlate of music processing as it occurs in a realistic or naturalistic environment, where eleven participants attentively listen to the whole piece of music (V. Alluri et al., 2012; I. Burunat et al., 2016). They successfully identified brain regions involved in processing of musical features in a naturalistic paradigm and found large-scale brain responses in cognitive, motor and limbic brain networks during continuous processing of low-level (timbral) and high-level (tonal and rhythmical) acoustic features using functional magnetic resonance imaging (fMRI). This was among the first approaches at attempting to discover the functional brain topology of musical processing by jointly drawing on computational feature extraction, behavioral, and brain-activity measures to isolate the variance associated with a number of musical features. Burunat et al. studied the replicability of Alluri’s findings using a similar methodological approach and a similar group of participants (Iballa Burunat et al., 2016). Unfortunately, all BOLD measurements by fMRI are to some degree confounded since they are indirect assessments of brain activity; they relate to blood flow and not to electrical processes and are therefore limited by poor temporal resolution due to the protracted hemodynamic response (Matthew J Brookes et al., 2014). Soon after, Cong et al used an analogous technique to investigate neural rhythms based on ongoing EEG data collected during same music stimuli (Cong, Alluri, Nandi, Toiviainen, Fa, Abu-Jamous, Gong, Craenen, Poikonen, & Huotilainen, 2013; Wang et al., 2016). They found the theta and alpha oscillations along central and occipital area of scalp topology seems significantly associated with high-level (tonal and rhythmical) acoustic features processing. During this period, many other studies tried to detect heard music from brain EEG signal using the time structure in music (Rebecca S Schaefer, Farquhar, Blokland, Sadakata, & Desain, 2011; Treder, Purwins, Miklody, Sturm, & Blankertz, 2014). Also, some researchers explored the brain responses to naturalistic music stimuli using EEG (Rogenmoser, Zollinger, Elmer, & Jäncke, 2016; Irene Sturm, Dähne, Blankertz, & Curio, 2015). Although, those sensor-level EEG studies analyzed the neural oscillations in human brain during music listening, the estimation of brain activity is performed at the sensor-level which impedes comparison with the rapidly growing literature on music-listening brain networks using fMRI. In addition, the interaction between spatial profiles and spectral bands is not yet fully understood.

In recent studies, resting state networks (RSNs) have been demonstrated by combining source localization with temporal/spatial ICA of the time frequency representation of MEG data (M. J. Brookes et al., 2011; Nugent et al., 2017; P. Ramkumar, Parkkonen, & Hyvarinen, 2014) and EEG data (Chen, Ros, & Gruzelier, 2013; Sockeel, Schwartz, Pelegrini-Issac, & Benali, 2016), in accordance with resting state fMRI networks. Additional studies have demonstrated that brain rhythms in a broad range of frequencies contributed to these networks. Brookes et al. examined that different spectral rhythms contributed differently to networks derived by individual ICA method (M. J. Brookes et al., 2012). For instance, beta and gamma oscillations contributed strongly to the bilateral insular networks, while a bilateral visual network was mainly driven by beta rhythms. Brookes et al. found that frequency-dependent network abnormalities may be associated with the pathophysiology of schizophrenia (M. J. Brookes et al., 2016). Hillebrand et al. found the frequency-specific patterns of information flow in RSNs, which shown direction and magnitude of information flow are different across frequencies (Hillebrand et al., 2016). Furthermore, Nugent et al. demonstrated differences in the resting state sensorimotor networks across frequency bands using a Multiband ICA technique(Nugent et al., 2017). However, these previous studies focus on networks under resting state, which is relatively simple to analyze. It is not clear whether the networks under real-world experience are frequency-specific.

Our hypothesis is that spatial profile under naturalistic stimuli differs across frequency bands as similarly as in resting states. In this paper, we propose a novel approach that combines music information retrieval (MIR) and spatial ICA to probe the spatial spectral patterns under music listening. EEG data were collected while participants were listening to a long piece of music. Short Time Fourier Transform was applied to EEG data. A three-way data was obtained. Source localization based on minimum-norm estimate was performed to get cortical source data. The cortical three-way data were rearranged into matrix. Then a complex-valued ICA was adopted to extract independent spatial spectral patterns and the time course of the patterns. Five long-term musical features were computationally extracted from the musical stimulus. Finally, the music-induced patterns were disentangled from the spontaneous activity by analysis of correlation between time courses of patterns and musical feature time series. Importantly, without any prior assumption, the frequency-dependent networks involved in musical feature processing were presented in a purely data-driven way.

## Material and methods

### Data acquisition

#### Participants

Fourteen right-handed and healthy adults aged 20 to 46 years old were recruited to take part in the current experiment after signing written informed consent. None of them was reported about hearing loss or history of neurological illnesses and none of them had musical expertise. This study was approved by the local ethics committee. All participants’ primary language was Finnish. More details on these participants can be found from our previous paper (Cong, Alluri, Nandi, Toiviainen, Fa, Abu-Jamous, Gong, Craenen, Poikonen, Huotilainen, et al., 2013).

#### EEG data acquisition

During the experiment, participants were informed to listen to the music with eyes opening. A 512s long musical piece of modern tango by Astor Piazzolla was used as the stimulus. Music was presented through audio headphones with about 30 dB of gradient noise attenuation. This music clip had appropriate duration for the experimental setting, because of its high range of variation in several musical features such as dynamics, timbre, tonality and rhythm (V. Alluri et al., 2012). The EEG data were recorded according to the international 10-20 system with BioSemi electrode caps (64 electrodes in the cap and 5 external electrodes at the tip of the nose, left and right mastoids and around the right eye both vertically and horizontally). EEG were sampled at a rate of 2048 Hz and stored for further processing in off-line. The external electrode at the tip of the nose was used as the reference. EEG channels were re-referenced using a common average. The data preprocessing was carried out using EEGLAB (Delorme & Makeig, 2004). The EEG data were visually inspected for artefacts and bad channels were interpolated using a spherical spline model. A notch filter at 50 Hz was applied to remove noise. High-pass and low-pass filter with 1 Hz and 30 Hz cutoff frequencies were then applied as our previous investigation of the frequency domain revealed that no useful information was found in higher frequencies (Cong, Alluri, Nandi, Toiviainen, Fa, Abu-Jamous, Gong, Craenen, Poikonen, & Huotilainen, 2013). Finally, the data were down-sampled to 256 Hz. In order to remove EOG (i.e., eye blinks), independent component analysis (ICA) was performed on EEG data of each participant. To additionally remove any DC-jumps occasionally present in the data, we differentiated each time series, applied a median filter to reject large discontinuities and reintegrated the signals back (Pavan Ramkumar, Parkkonen, Hari, & Hyvärinen, 2012).

#### Musical features

Based on the length of the window used in the computational analyses, the musical features can be generally classified into two categories: long-term features and short-term features (V. Alluri et al., 2012; Cong, Alluri, Nandi, Toiviainen, Fa, Abu-Jamous, Gong, Craenen, Poikonen, Huotilainen, et al., 2013). Five long-term musical features including Mode, Key Clarity, Fluctuation Centroid, Fluctuation Entropy and Pulse Clarity were examined here. They were extracted using a frame-by-frame analysis approach commonly used in the field of Music Information Retrieval (MIR). The duration of the frames was 3s and the overlap between two adjacent frames 67% of the frame length. The chosen length of the frame was approximately consistent with the length of the auditory sensory memory (V. Alluri et al., 2012). This analysis process yielded the time series of musical feature at a sampling frequency of 1 Hz, in accordance with the short-time Fourier transform (STFT) analysis of EEG data. Thus, both the musical features and temporal courses of EEG had 512 time points. All the features were extracted using the MIRtoolbox (Lartillot, Toiviainen, & Eerola, 2008) in MATLAB environment.

For the completeness of the content, we briefly introduce the five features below. We extracted two tonal and three rhythmic features. For the tonal features, Mode represents the strength of major or minor mode. Key Clarity is defined as the measure of the tonal clarity. The rhythmic features included Fluctuation Centroid, Fluctuation Entropy and Pulse Clarity. Fluctuation Centroid is the geometric mean of the fluctuation spectrum, representing the global repartition of rhythm periodicities within the range of 0-10Hz (V. Alluri et al., 2012). This feature indicates the average frequency of these periodicities. Fluctuation entropy is the Shannon entropy of the fluctuation spectrum, representing the global repartition of rhythm periodicities. Fluctuation entropy is a measure of the noisiness of the fluctuation spectrum (V. Alluri et al., 2012; Cong, Alluri, Nandi, Toiviainen, Fa, Abu-Jamous, Gong, Craenen, Poikonen, Huotilainen, et al., 2013). Pulse Clarity, naturally, is an estimate of clarity of the pulse (V. Alluri et al., 2012; Cong, Alluri, Nandi, Toiviainen, Fa, Abu-Jamous, Gong, Craenen, Poikonen, Huotilainen, et al., 2013).

### Source Localization

For each subject, the brain’s cortical surface was reconstructed from an anatomical MRI template in Brainstorm (Tadel, Baillet, Mosher, Pantazis, & Leahy, 2011). Dipolar current sources were estimated at cortical-constrained discrete locations (source points) separated by 15 mm. Each hemisphere was modelled by a surface of approximately 2,000 vertices, thus a mesh of approximately 4,000 vertices modelled the cortical surface for each subject.

The measured EEG signals are generated by postsynaptic activity of ensembles of cortical pyramidal neurons of the cerebral cortex. These cortical pyramidal neurons can be modelled as current dipoles located at cortical surface (Lin, Belliveau, Dale, & Hamalainen, 2006). The scalp potentials generated by each dipole depend on the characteristics of the various tissues of the head and are measured by the EEG scalp electrodes. With the geometry of the anatomy and the conductivity of the subject’s head, the time course of the dipole’s activity can be assessed by solving two consecutive problems: the forward problem and the inverse problem.

The forward problem is to model the contribution of each dipole to the signals of the EEG electrodes by solving Maxwell’s equations, which takes the geometry and conductivity of head tissues into account. In this study, a forward solution was computed calculated using the symmetric boundary element method (BEM) for each source point while a relative conductivity coefficient was assigned to each tissue (with default MNI MRI template).

To solve the inverse problem, minimum-norm estimate (Lin et al., 2006) was adapted with a loose orientation constraint favoring source currents perpendicular to the local cortical surface (no noise modelling). When computing the inverse operator (1) the source orientations were constrained to be normal to the cortical surface; (2) a depth weighting algorithm was used to compensate for any bias affecting the superficial sources calculation; and (3) a regularization parameter, λ^2^ = 0.1 was used to minimize numerical instability, and to effectively obtain a spatially smoothed solution. Finally, an inverse operator G of dimensions *N*_*s*_ × *N*_*c*_ (where *N*_*s*_ is the number of source points and *N*_*c*_ is the number of channels:*N*_*s*_ ≫ *N*_*c*_) was obtained to map the data from sensor-space to source-space. Here, we had *N*_*s*_ = 4000 and *N*_*c*_ = 64.

### Spatial Fourier Independent Component Analysis

#### Time-frequency data in cortical source space

Preprocessed EEG data *Y*_0_ (*N*_*c*_ channels x *N*_*p*_ sampling points) were transformed by short-time Fourier transform (STFT) to obtain complex-valued time-frequency representation (TFR) data *Y*_1_ (*N*_*c*_, *N*_*f*_, *N*_*t*_). To obtain TFR data in source space, three-way sensor-space TFR data *Y*_1_ was reorganized as two-way matrix 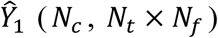. The source-space TFR data 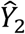 was then obtained by left-multiplying the linear inverse operator G (*N*_*s*_, *N*_*c*_) which was computed using minimum-norm estimate inverse solution sensor-space data 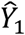,

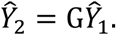

Two-way data 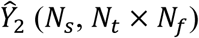 can be rearranged as a three-way tensor format *Y*_2_ (*N*_*s*_, *N*_*t*_, *N*_*f*_). For application of spatial Fourier ICA, we then rearranged the three-way tensor *Y*_2_ as a two-way matrix X_0_ (*N*_*t*_,*N*_*f*_ x *N*_*s*_). Thus, each row of *X*_0_ was comprised of the complex-valued short-time Fourier coefficients from each source point for specific time points and each column represented a time point corresponding to a short-time window. In this study, the Hamming-widow with 3-second-length and 2-second-overlap of the adjacent windows was selected, resulting in a sampling rate of 1 Hz in time dimension. This sampling rate was in consistent with musical feature time series (see Musical features). The duration of EEG was 512s, so we had *N*_*t*_ = 512 time points. We adopted a 512-point FFT to calculate the STFT resulting in 256 frequency bins (Range of frequency: 1-128 Hz) for each window. We selected the range of frequency bins covering 1-30 Hz (*N*_*f*_ = 60) for further analysis.

#### Application of complex-valued ICA on reshaped data

For data *X*_0_, we applied complex-valued ICA (A. Hyvarinen, Ramkumar, Parkkonen, & Hari, 2010) and treated each row as an observed signal assumed to be a linear mixture of unknown spatial spectral pattern. Since the original data (*X*_0_) dimension was relatively high for the complex ICA calculation, data dimension reduction was required in the preprocessing step of ICA. A common approach of data dimension reduction is principal component analysis (PCA) which is linear. Here we extended PCA to the complex domain by considering complex-valued eigenvalue decomposition (Li, Correa, Rodriguez, Calhoun, & Adali, 2011). The choice of model order was based on previous studies (Abou - Elseoud et al., 2010; Smith et al., 2009), which suggested the number of a dimension slightly larger than the expected number of underlying sources. In this study, we tried different model orders and found that 20 was a reasonable order, which preserved much of the information in the data and reduced the dimensionality of the results. Then we extracted 20 independent components using complex-valued FastICA algorithm which applied ICA to STFT of EEG data in order to find more interesting sources than with time-domain ICA (A. Hyvarinen et al., 2010). This method is especially useful for finding sources of rhythmic activity. After complex-valued ICA, a mixing matrix 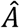 (*N*_*t*_, *N_ic_* = 20 is number of components) and estimated source matrix 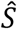 were obtained. Each column of 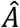 represented the temporal course for each independent component (IC). The ICs in the rows of 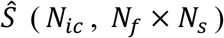 represented spatial spectral patterns, which can be decomposed into the spatial power map and power spectra.

#### Spatial map, spectrum and temporal course of ICs

By reshaping each row of 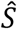 for each IC, we obtained a matrix (*N*_*f*_, *N*_*s*_), which meant there was a Fourier coefficient spectrum for each cortical source point. To obtain and visualize the spatial map of the IC, we computed the average of the squared magnitude of the complex Fourier coefficients across those frequency bins. Since the distribution of mean squared Fourier amplitude over the whole brain is highly non-Gaussian, we did not apply conventional z-score-based thresholding; instead, we applied a threshold to display for each component map only source points with the top 5% squared Fourier amplitude (Pavan Ramkumar et al., 2012). Then we analyzed the correlation coefficient of the spatial maps in those frequency bins and those spatial maps were similar. To visualize and obtain the spectrum of each IC, we calculated the mean of the Fourier power spectrum across those source points exceeding the 95^th^ percentile (Pavan Ramkumar et al., 2012). Finally, we extracted the absolute values of the column of mixing matrix *Â* corresponding to the row of the estimated IC as the time course, which reflected fluctuations of the Fourier amplitude envelope for the specific frequency and spatial profile.

### Stability of ICA decomposition

To examine the stability of ICA, we applied 100 times ICA decomposition for each subject with different initial conditions. For the real-valued case, ICASSO toolbox (Himberg, Hyvarinen, & Esposito, 2004) has been used to evaluate stability among multiple estimates of the fastICA algorithm (Aapo Hyvarinen, 1999). All the components estimated from all runs were collected and clustered based on the absolute value of the correlation coefficients among the squared source estimates of ICASSO. Finally, the stability index Iq was computed for each component. Iq reflects the isolation and compactness of a cluster (Himberg et al., 2004). Iq is calculated as follows:

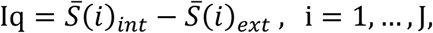

where 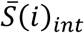 denotes the average intra-cluster similarity; 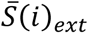 indicates average inter-cluster similarity and J is the number of clusters. The Iq ranges from ‘0’ to ‘1’. When Iq approaches ‘1’, it means that the corresponding component is extracted in almost every ICA decomposition application. This indicates a high stability of the ICA decomposition for that component. Otherwise, it means the ICA decomposition is not stable. Correspondingly, if all the clusters are isolated with each other, ICA decomposition should be stable. In general, there is no established criterion upon which to base a threshold for cluster quality. Given the preliminary nature of this investigation, we consider the decomposition is stable if the Iq is greater than 0.65.

In this study, the ICASSO toolbox was modified to be available for the complex-valued case as well. The correlation matrix was used as the similarity measure for clustering in real-valued ICASSO. For the complex case, since the ICs were complex-valued, we just considered the correlation matrix among the magnitude ICs to perform the clustering (Li et al., 2011). Then, we took the Iq as the criterion to examine stability of the ICA estimate.

**Fig. 1:**
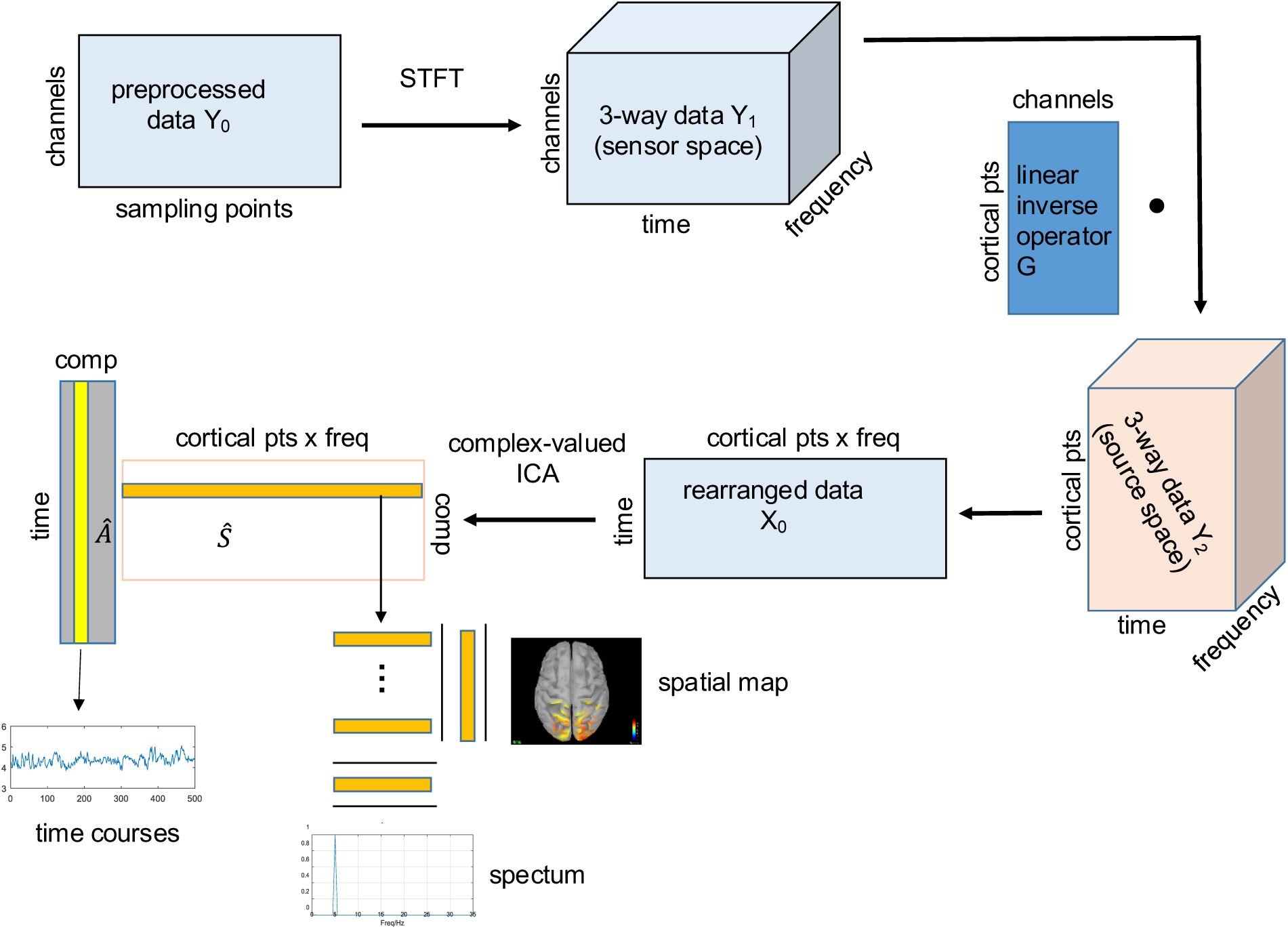
Diagram of steps for spatial ICA. Preprocessed EEG data ***Y***_**0**_ (*N*_*c*_ channels x *N*_*p*_ sampling points) was transformed by short-time Fourier transform (STFT) to obtain time-frequency representation (TFR) data ***Y***_**1**_ (*N*_*c*_, *N*_*f*_, *N*_*t*_). Source-space data ***Y***_**2**_ (*N*_*s*_,*N*_*f*_,*N*_*t*_) were obtained by left-multiplying inverse operator **G** to sensor-space data ***Y***_**1**_.Reshape three-way data ***Y***_**2**_ to matrix data ***X***_**0**_ (*N*_*t*_, *N_s_* x *N*_*f*_). Complex-valued ICA was done on this matrix. The squared absolute values of the columns of mixing matrix provide the time course of Fourier-power and the rows of source matrix were decomposed into the spatial power map and spectrum (see text).

### Testing for stimulus-related networks

After ICA decomposition, we obtained 20*14=280 ICs (14 subjects, 20 components for each subject). Now the challenge is to determine which one of these represents the genuine brain responses. In all ICA based methods, it is a general question that which independent components need to be retained or which component just reflects noise. Here, we examine which components were modulated significantly by the musical features. We computed the correlation (Pearson’s correlation coefficient) between the time courses of musical features and the time courses of those ICs (the dimensionality of both them is 512 points) in order to select stimulus-related activations. We used the Monte Carlo method and permutation tests presented in our previous research (V. Alluri et al., 2012; Cong, Alluri, Nandi, Toiviainen, Fa, Abu-Jamous, Gong, Craenen, Poikonen, Huotilainen, et al., 2013) to calculate the threshold of significant correlation coefficient. In this method, a Monte Carlo simulation of the approach was performed to determine the threshold for multiple comparisons. We kept those ICs whose time courses were significantly correlated (p<0.05) with the time courses of musical features for further analysis.

### Cluster analysis

The selected ICs had been represented by spatial map, spectrum and temporal course. Since spatial ICA was carried out on individual level EEG data, we needed to examine the inter-subject consistency among participants. In this study, we focused on the spatial pattern emerging in the process of freely listening to music, so a group level data analysis was performed by clustering spatial maps of the selected ICs to evaluate the consistency among the participants. For reliable clustering, we applied a conventional z-score-based normalization to each spatial map. All spatial maps of the screened components significantly correlated with musical features were clustered into M clusters to find common spatial patterns among most of participants. Here for simplicity, a conventional k-means cluster algorithm was used with the Kaufman Approach (KA) for initializing the algorithm. We used the minimum description length (MDL) to determine the number of clusters M. Afterwards we countered the number of subjects involved in ICs in each cluster. If the number of subjects in one cluster is less than half of the all subjects, this cluster would be discarded for the reason that such a cluster does not reveal information shared among enough participants. For the retained clusters, the spatial-spectral-temporal information was obtained, which was represented by the centroid of the cluster, the spectra of ICs and the numbers of subjects whose temporal courses were involved in this cluster. Figure 2 shows the flow of group level analysis.

**Fig. 2.**
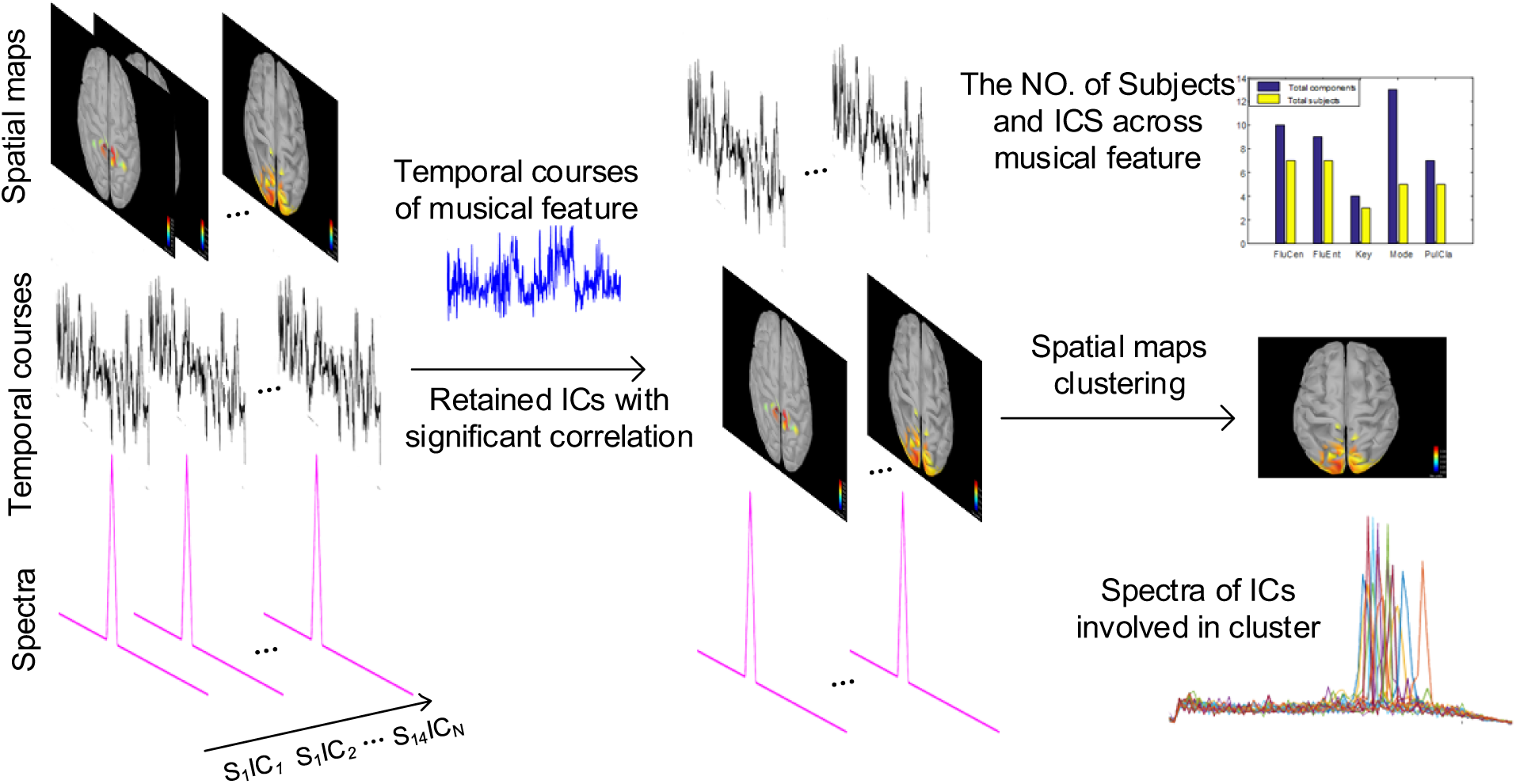
Diagram of cluster analysis. Left column: The spatial maps, time courses and spectra of extracted IC from all subjects (S_M_IC_N_ denotes number #N components from subjects #M). Middle column: One example cluster. The spatial maps, time courses and spectra of retained ICs after correlation analysis with musical feature. Right column: Clustered spatial map, corresponding spectrum, and the distribution across musical feature.

## Results

### Musical Features

Five musical features were extracted by MIRtoolbox (Olivier Lartillot, 2007) with 3s time-widow and 2s overlap, resulting in 1 Hz sampling rate of temporal course. They are Fluctuation Centroid, Fluctuation Entropy, Key Clarity, Mode and Pulse Clarity. The time series of these features had a length of 512 samples, which matched the length of the time course of the EEG components. Figure 3 shows their temporal courses.

**Fig. 3.**
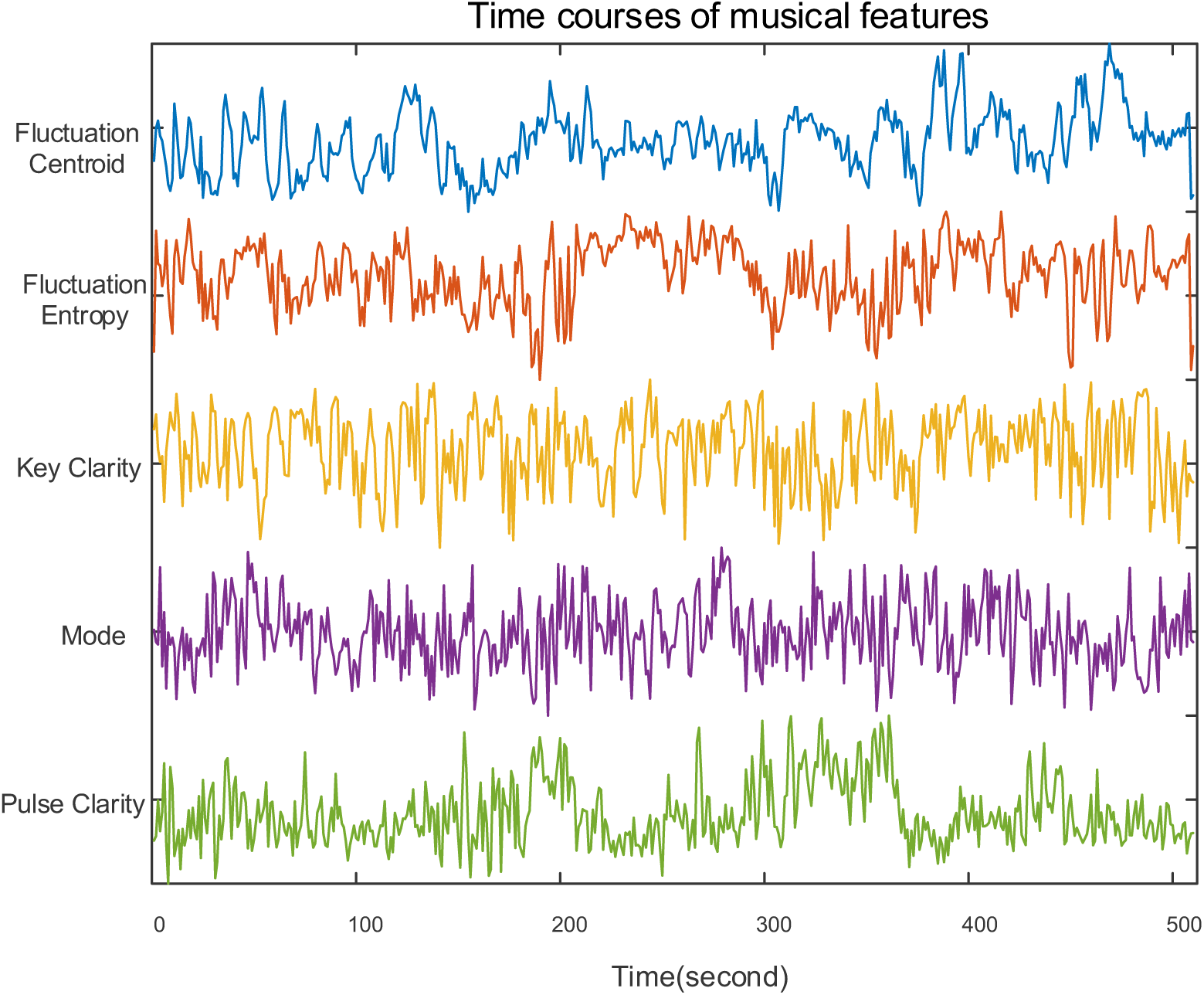
Time course of five musical features

### Stability of ICA decomposition

We extracted 20 ICs using modified ICASSO with 100 runs for each subjects’ data, then we obtained the stability index Iq. Figure 4 shows the magnitude of Iqs for each participant, greater than 0.7 for most ICs. The 20 ICs were separated with each other for every participant from the view of clustering. Thus, the ICA estimate was stable and the results of ICA decomposition in this study were satisfactory for each participant data to further analysis.

**Fig. 4.**
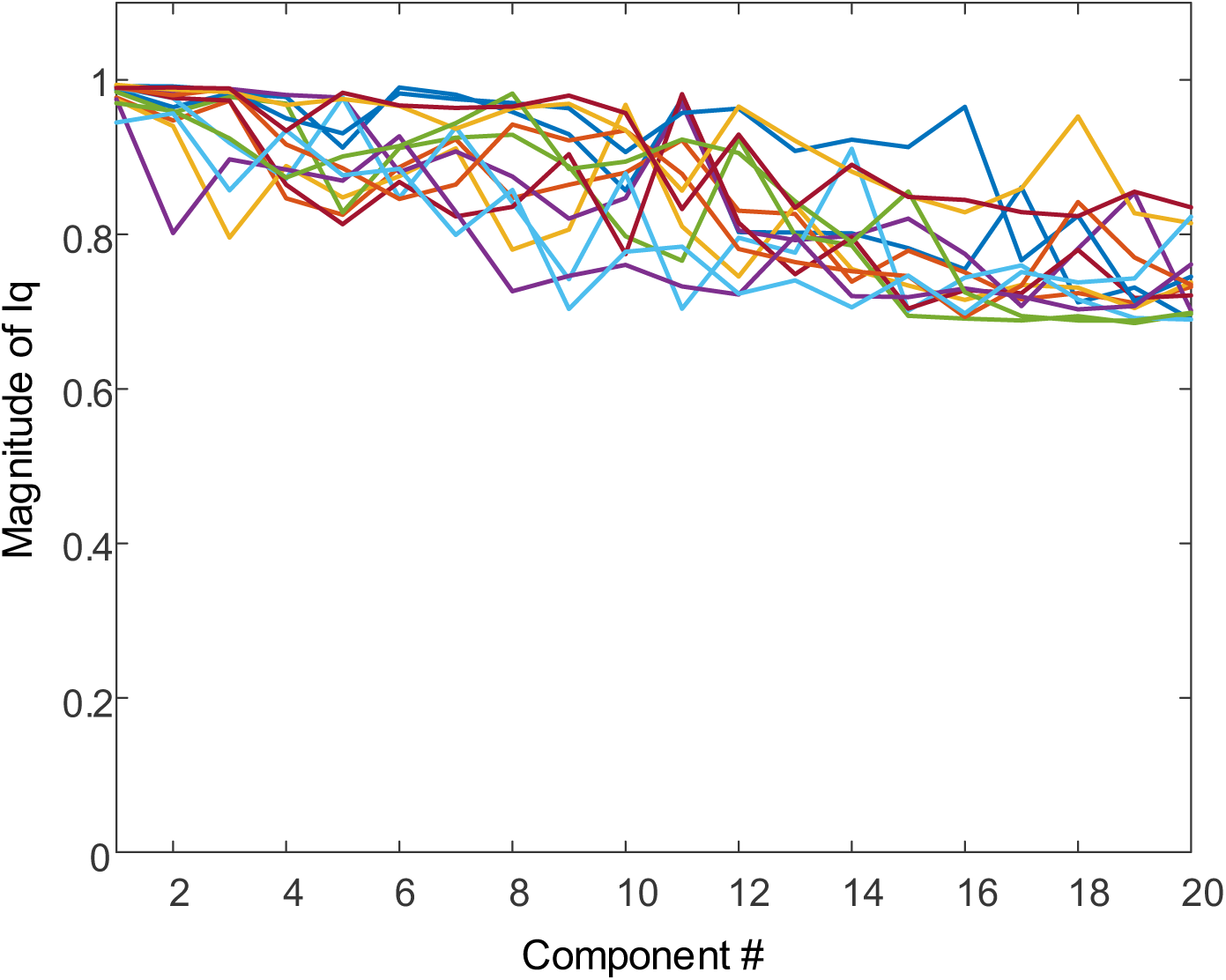
Iq of each component extracted. Different curves represent different participants

### Interesting clusters: frequency-specific networks

After 85 ICs whose spatial maps were significantly correlated with musical features were selected, we set the number of clusters as five by performing MDL to estimate the optimal model order. Then the spatial maps of ICs were clustered into five clusters. Three clusters representing frequency-specific networks were chosen since the number of subjects in the cluster is more than half of the all subjects. Figure 5 demonstrates one of these clusters including the centroid of all spatial maps (Fig. 5A), the distribution of number of subjects across musical features (Fig. 5B) and the spectrum of the ICs in this cluster (Fig. 5C). Then we computed the correlation coefficients among spatial maps in each cluster to evaluate the performance of clustering. Figure 8 shows the inter-cluster similarity. We computed the mean of the correlation coefficients in each cluster and the corresponding standard deviation (SD). For cluster#1, the mean is 0.642 and the corresponding standard deviation (SD) is 0.1238. For cluster#2, the mean is 0.7125 and SD is 0.0572. For cluster#3, the mean is 0.8084 and SD is 0.0747. This indicates that the spatial patterns are similar across the participants. In the Table 1, we listed the participants whose EEG data were correlated with every musical feature in each cluster.

**Table 1.** Participants involved in each cluster across musical features among 14 subjects (from 1 to 14)

| Cluster | Musical features |  |  |  |  | Total subjects |
| --- | --- | --- | --- | --- | --- | --- |
|  | Fluctuation<br>Centroid | Fluctuation<br>Entropy | Key Clarity | Mode | Pulse Clarity |  |
| #1 | 2 4 5 12 | 2 | 1 2 3 13 | 4 | 5 | 7 |
| #2 | 1 8 10 14 | 8 11 13 | 7 8 14 | 2 8 11 14 | 1 10 13 | 8 |
| #3 | 4 5 7 9 11 | 3 6 | 5 6 9 | 2 3 4 6 7 9 | 7 | 8 |

**Fig. 5.**
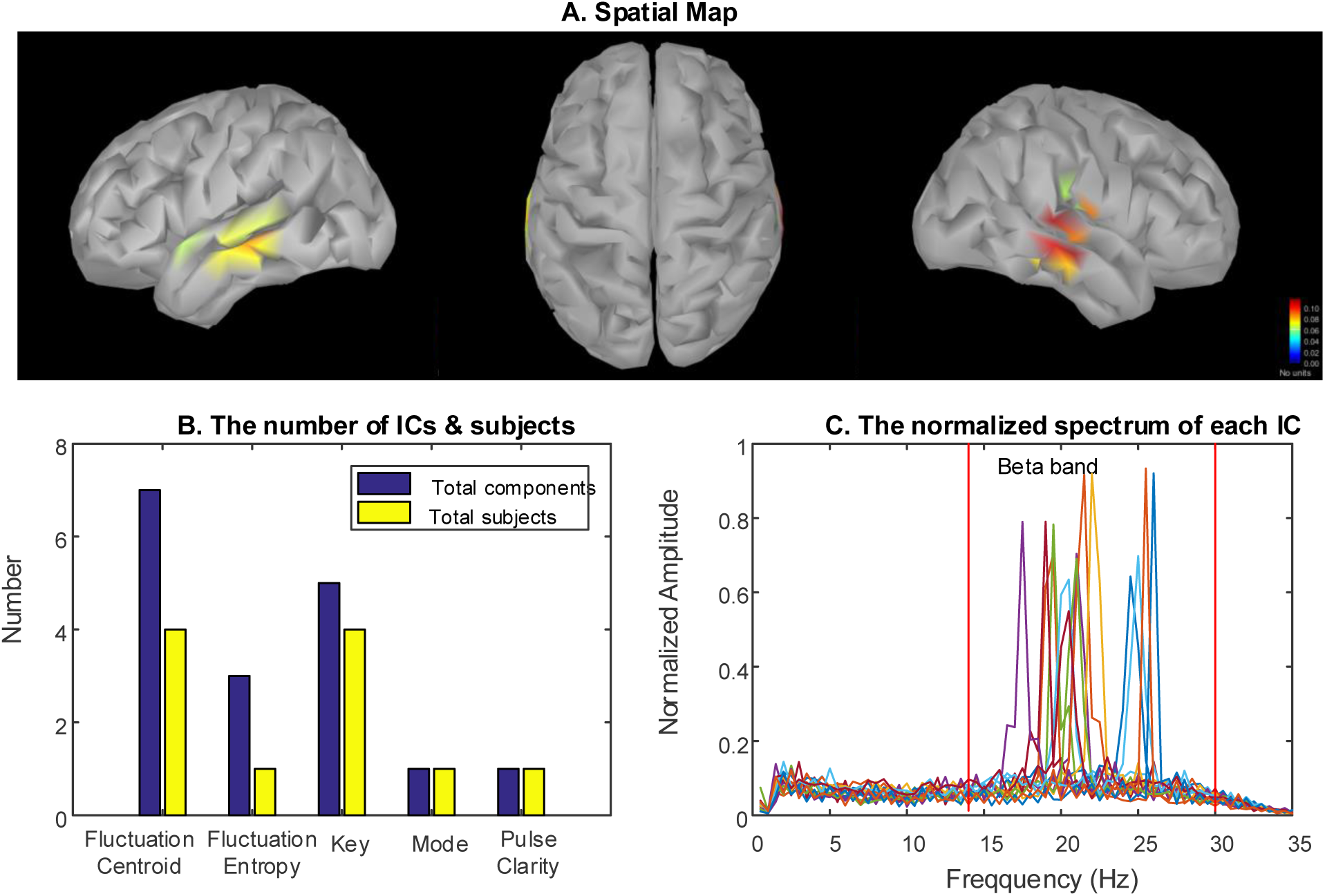
Cluster#1: Beta-specific networks. A) The centroid of the spatial maps in the cluster, which reflected a spatial pattern across most of participants. The bilateral superior temporal gyrus (STG) was activated (from left to right: left hemisphere, top view, right hemisphere). B) The number of ICs and subjects involved in the cluster were distributed across musical features. C) The spectra of each components were located in delta or beta band. Different curves represent different ICs.

#### Beta-specific network

Figure 5 shows results of the Beta-specific brain networks engaged in processing music features. The spatial map displays that musical features were associated with increased activation in the bilateral superior temporal gyrus (STG). The spectrum of ICs in this cluster illustrates the beta rhythm (focusing on 20 Hz) was involved in generating this network. Thus, relatively large-scale brain region generated by beta rhythm was activated in the bilateral STG and the magnitude of activation in right hemisphere was a little stronger than left hemisphere. This Beta-specific network was found in seven subjects during music free-listening (see the first row of Table 1). Fluctuation Centroid were associated with this brain networks among subjects 2, 4, 5, and 12. The brain networks of subjects 1, 2, 3 and 13 were correlated with Key feature. For Fluctuation Entropy, Pulse Clarity and Mode, there was one subject involved in this cluster respectively. In addition, the number of ICs correlated with the musical features was more than the number of participants since there were 20 ICs for each subject.

#### Alpha-specific network

Figure 6 displays relatively large brain activity in the bilateral occipital lobe according to the spatial map. As can be seen, the oscillations of this pattern were dominated by alpha rhythm (focusing on 10 Hz) with few ICs located in Delta band. There were eight participants appearing alpha-specific occipital networks under free-listening to music. The second row of the Table 1 shows the subjects involved in the networks linked with each musical feature.

**Fig. 6.**
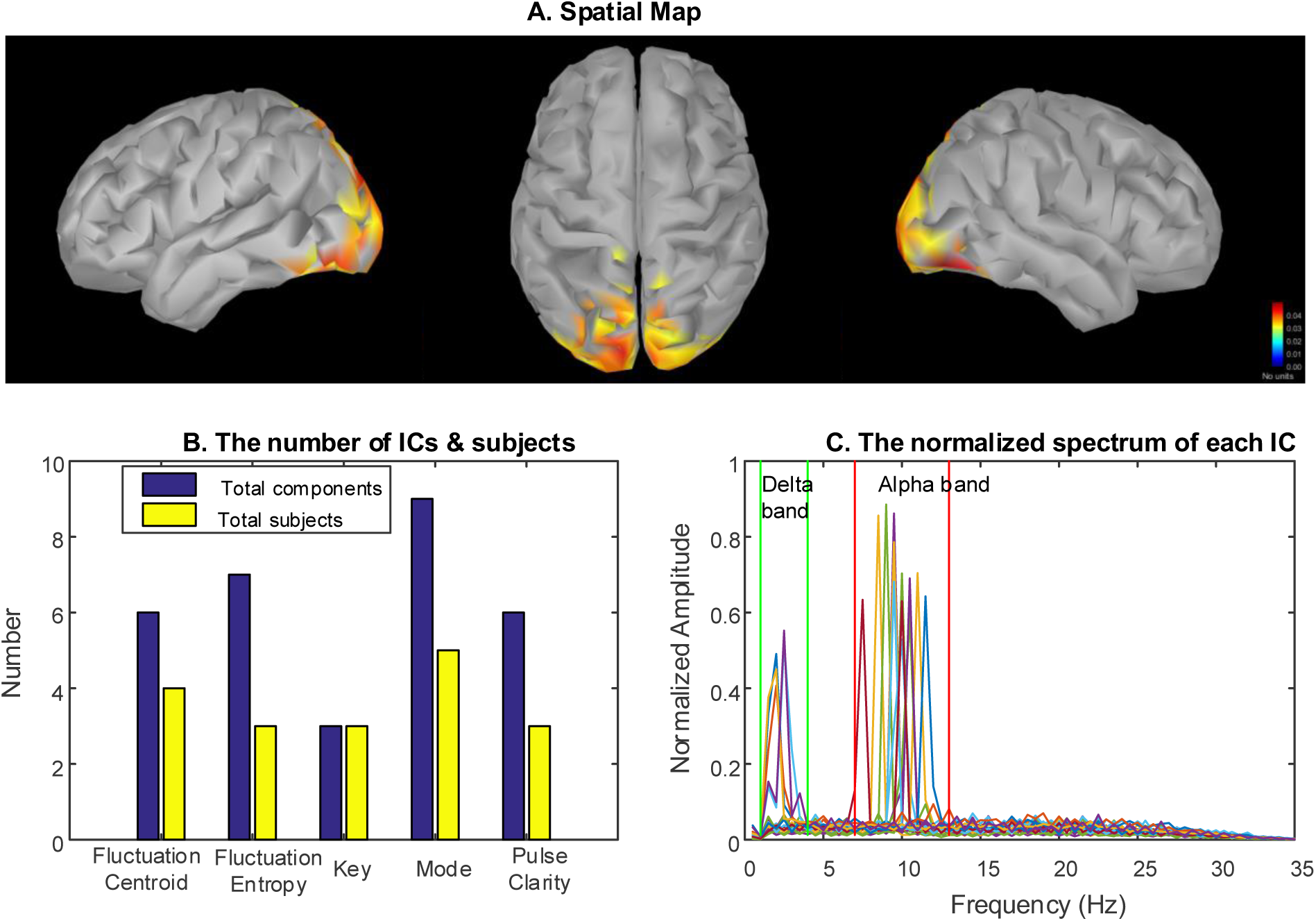
Cluster#2: Alpha-Delta-specific networks. A) The centroid of the spatial maps in the cluster, which reflected a spatial pattern across most of participants. As can be seen that the bilateral occipital cortex was activated (from left to right: left hemisphere, top view, right hemisphere). B) The number of ICs and subjects involved in the cluster were distributed across musical features. C) The spectra of each components were located in delta or beta band. Different curves represent different ICs.

#### Delta-Beta-specific network

Figure 7A illustrates increased activity linked with musical features in bilateral prefrontal gyrus (PFG). The spectrum (Fig. 7C) shows both beta and delta oscillations recruited these areas across participants. The delta-beta-specific networks were found in eight subjects. Mode was associated with this brain networks among subjects 2, 3, 4, 6, 7 and 9. The networks of subjects 4, 5, 7, 9 and 11 were correlated with Fluctuation Centroid (see the third row of Table 1).

**Fig. 7.**
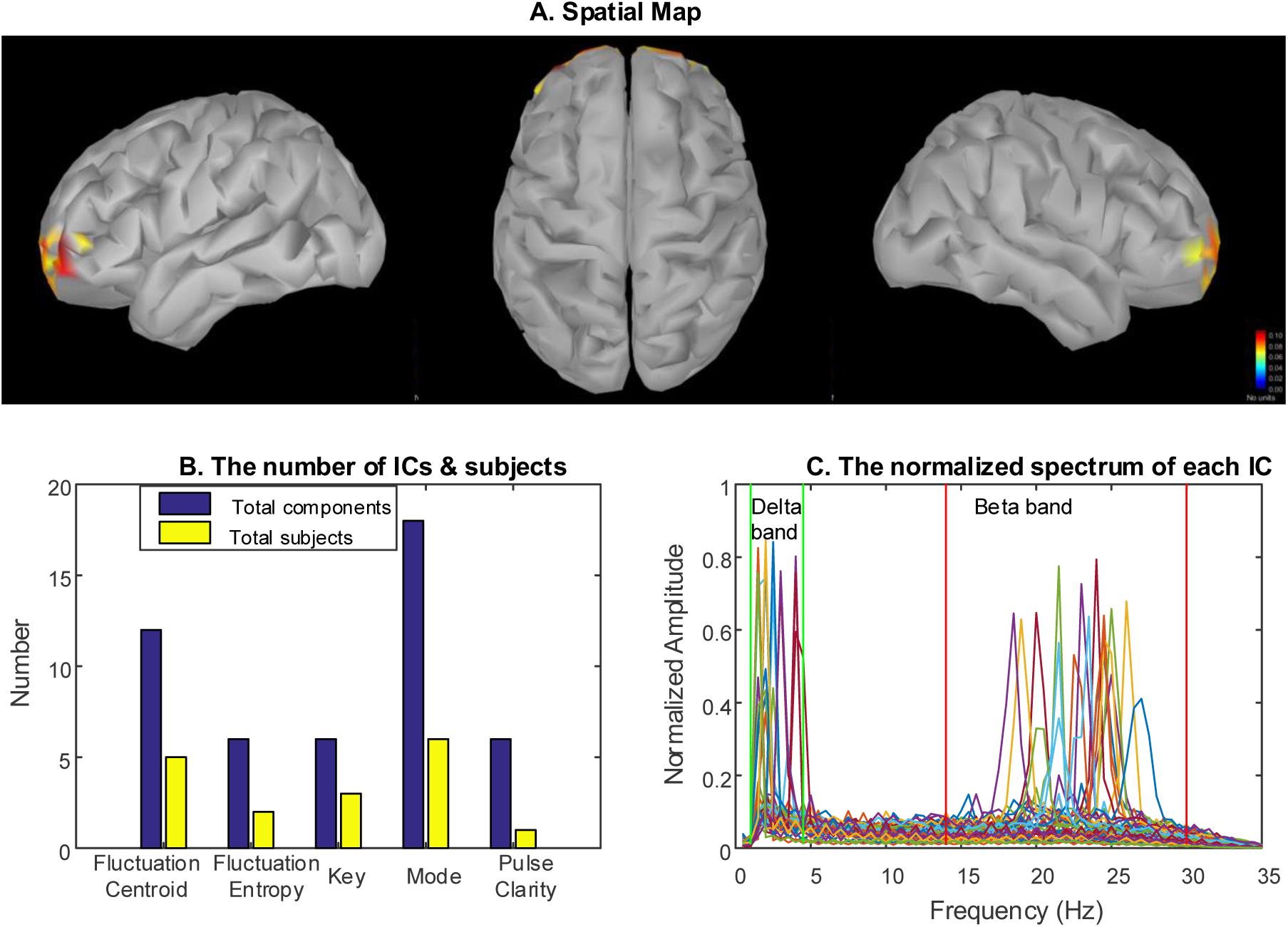
Cluster#3: Alpha-Beta-specific networks. A) The centroid of the spatial maps in the cluster, which reflected a spatial pattern across most of participants. As can be seen that the bilateral prefrontal cortex was activated (from left to right: left hemisphere, top view, right hemisphere). B) The number of ICs and subjects involved in the cluster was distributed across musical features. C) The spectra of each components were located in delta or beta band. Different curves represent different ICs.

**Fig. 8.**
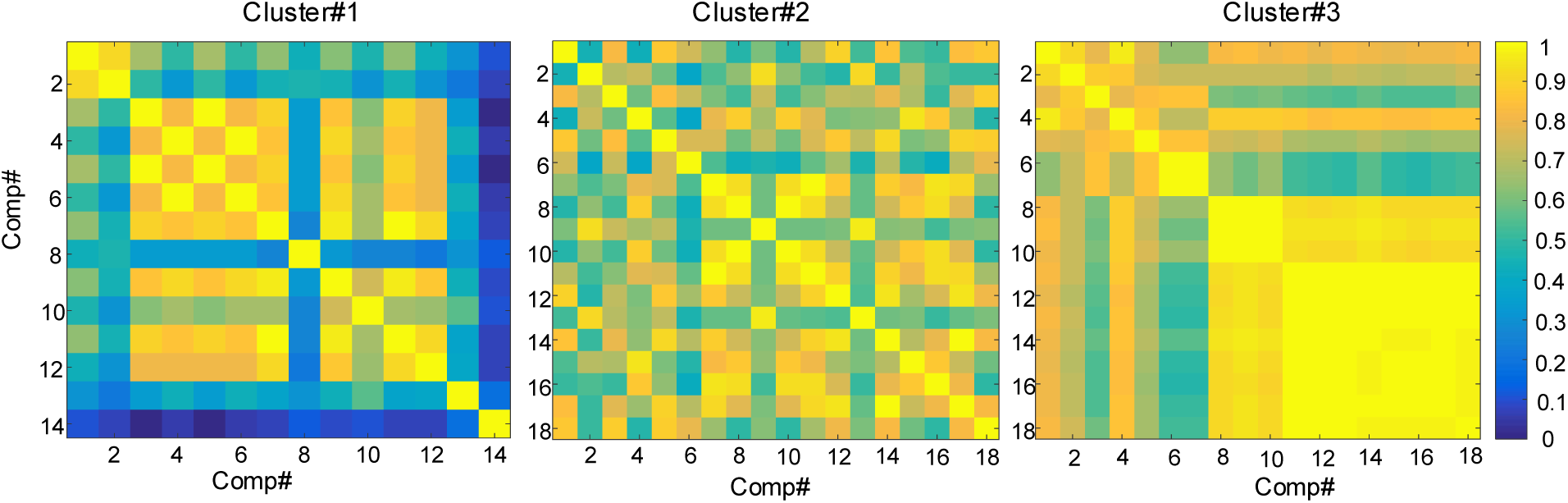
Correlation coefficients matrix among spatial maps of the ICs in each cluster. The mean correlation coefficient in cluster#1 is 0.642 and the corresponding standard deviation (SD) is 0.1238. For cluster#2 the mean is 0.7125 and SD is 0.0572. For cluster#3, the mean is 0.8084 and SD is 0.0747.

## Discussion

In this study, we investigated spatial spectral profiles of brain networks during music free-listening. To this end, we proposed a novel method combing spatial ICA, source localization and music information retrieval. EEG data were recorded when participants listened to a piece of music freely. First, we applied short-time Fourier transform to preprocessed EEG data. After this, an inverse operator was obtained using source localization and the sensor-space data was mapped to source-space data. Then complex-valued ICA was performed to extract spatial spectral patterns. The stability of ICA estimate was evaluated using a complex-value ICASSO. Meanwhile, the temporal evolutions of five long-term musical features were extracted by the commonly used MIRtoolbox. Following this, the spatial spectral ICs were chosen by correlating their temporal course with the temporal course of musical features. To examine the inter-subject consistency, a cluster analysis was applied to spatial patterns of the retained ICs. Overall, our results highlighted the frequency-dependent brain networks during listening music freely. The results are consistent with previous findings published in other studies (V. Alluri et al., 2012; Cong, Alluri, Nandi, Toiviainen, Fa, Abu-Jamous, Gong, Craenen, Poikonen, Huotilainen, et al., 2013; Janata et al., 2002).

It was found that beta-specific brain networks in the bilateral STG emerged from dynamic processing of musical features (see Fig. 5). The bilateral STG were mostly activated during music listening, which was involved in long-term musical features processing. It was interesting to note that the beta oscillations were enhanced in this bilateral spatial profile (see Fig. 5C). This spatial spectral pattern appeared more related with Fluctuation Centroid and Key processing than Fluctuation Entropy, Mode and Pulse Clarity (see Fig. 5B). The same areas were found in previous studies where timbre-related features were correlated with activations in large areas of the temporal lobe using fMRI (V. Alluri et al., 2012). Besides, early MEG studies demonstrated that cortical rhythm activity in beta band activity (15-30Hz) is tightly coupled to behavioral performance in musical listening and associated with predicting the upcoming note events (Doelling & Poeppel, 2015). Our findings indicate that the specific beta rhythms appearing in the STG play an important function in music perception, which means the interplay between beta oscillations with the STG emerges during listening to a continuous and naturalistic music. We also observed alpha oscillatory visual networks (see Fig. 6), which is in line with our previous study (Cong, Alluri, Nandi, Toiviainen, Fa, Abu-Jamous, Gong, Craenen, Poikonen, Huotilainen, et al., 2013). Alpha-band oscillations play an important role in basic cognitive process, which is linked to suppression and selection of attention (Klimesch, 2012). The delta rhythms of few subjects had enhancement in this visual network, which is not yet understood. The alpha-specific power over visual cortices was larger when attention was focused on the auditory stimuli which is in line with those discussed by (Barbara F. Händel, 2011). A delta-beta oscillatory network in prefrontal cortex were also observed during listening to music (see Fig. 7). Helfrich et al. argued that the prefrontal cortex provides the structural basis for numerous higher cognitive functions and oscillatory dynamics of prefrontal cortex provide a functional basis for flexible cognitive control of goal-directed behavior (Helfrich & Knight, 2016). Besides, prefrontal cortex has the function of entrainment as a mechanism of top-down control (Helfrich & Knight, 2016). Our findings provided the evidence that the higher cognitive function were involved in continuous and naturalistic music. Janata et al. identified an area in the rostromedial prefrontal cortex as a possible brain units for tonal processing (Janata et al., 2002). In addition, some studies demonstrated that oscillations in the delta and beta bands were instrumental in predicting the occurrence of auditory targets (Arnal, Doelling, & Poeppel, 2015; Doelling & Poeppel, 2015). That may explain why the delta-beta oscillations in this study appears in prefrontal cortex.

However, most of these studies investigated one pattern of the spatial spectral profile and did not examined the interplay between spatial brain networks and spectral profile. In contrast, we studied the interactions between brain region and cortical oscillations and found the brain networks during music listening were frequency-specific. In terms of our proposed approach for analysis of frequency-specific networks during naturalistic music listening, we can credibly find the spatial spectral patterns elicited by musical stimulus. There are some related approaches using spatial ICA in a variety of specific techniques to investigate the RSNs under MEG data. Nugent et al. proposed a method named as MultibandICA to derive frequency-specific spatial profile in RSNs; Hilbert transform was first applied to MEG data to extract six frequency bands (delta, theta, alpha, beta, gamma, high gamma) and the six bands data were concatenated in certain dimensionality; ICA was then performed to concatenated data (Nugent et al., 2017). Similar methods were proposed in (Sockeel et al., 2016). Besides, there are some other spatial ICA approaches including spatial Fourier ICA (SFICA) (P. Ramkumar, Parkkonen, Hari, & Hyvarinen, 2012) and envelope Fourier ICA (eFICA) (P. Ramkumar et al., 2014). Those methods did not analyze the stability of ICA decomposition and they just considered the resting state without analyzing external stimulus features. Another important asset of our study is that the clustering was applied to the spatial maps to examine the inter-subject consistency in proposed method. The correlation coefficients were then computed in each cluster. We observed that the individual spatial spectral profiles in every retained cluster were similar but the corresponding time courses were different. This is different from analysis of event-related potential (ERP) where temporal ICA components sharing identical spatial profiles might be similar. The differences might be resulted from different responses of participants under real-word experiences. In the future, we will attempt to develop group spatial ICA to analyze group-level data where the individual data are concatenated in time dimension.

## Conclusion

In this study, we introduced a novel approach for exploiting the spectral-spatial structure of brain during naturalistic stimulus. A complex-value ICA applied to source-space time-frequency representation of EEG data. Following this, a modified ICASSO was performed to evaluate the stability of ICA estimate and a cluster analysis was applied to examine the inter-subject consistency. The identified networks involved in music perception were in line with those previous studies. Further, we found that brain networks under music listening were frequency-specific and three frequency-dependent networks associated with processing musical features were observed.

## Acknowledgment

This work was supported by the National Natural Science Foundation of China (Grant No. 81471742), the Fundamental Research Funds for the Central Universities [DUT16JJ(G)03] in Dalian University of Technology in China, and the scholarship from China Scholarship Council (No. 201600090042).

## Appendix A

The features were extracted from the stimulus on a frame-by-frame basis (see (Vinoo Alluri & Toiviainen, 2010) for more details). A brief description of each of the acoustic features is presented below. A detailed explanation can be found in the user manual of the MIRToolbox (Lartillot & Toiviainen, 2007).

Mode: strength of major of minor mode.

Key Clarity: the strength of the estimated key, computed as the maximum of cross-correlations between the chromagram extracted from the music and tonality profiles representing all the possible key candidates.

Fluctuation centroid: geometric mean of the fluctuation spectrum representing the global repartition of rhythm periodicities within the range of 0-10 Hz, indicating the average frequency of these periodicities.

Fluctuation entropy: Shannon entropy of the fluctuation spectrum (Pampalk et al., 2002) representing the global repartition of rhythm periodicities. Fluctuation entropy is a measure of the noisiness of the fluctuation spectrum. For example, a noisy fluctuation spectrum can be indicative of several co-existing rhythms of different periodicities, thereby indicating a high level of rhythmic complexity.

Pulse Clarity: the strength of rhythmic periodicities sound, representing how easily the underlying pulsation in music can be perceived.

